# *Brucella* species circulating in wildlife in Serengeti ecosystem, Tanzania

**DOI:** 10.1101/2020.02.06.936807

**Authors:** R. M. Sambu, C. Mathew, H.E. Nonga, A. S. Lukambagire, Richard B. Yapi, G. Fokou, J. Keyyu, B. Bonfoh, R. R Kazwala

## Abstract

**Background:** Brucellosis is a bacterial zoonosis of public health and economic importance world-wide. It affects a number of domestic animals, wildlife and humans. This study was carried out to determine circulating *Brucella* species in wildlife in Serengeti ecosystem using molecular techniques.

**Methodology:** A total of 189 samples including EDTA blood, serum and amniotic fluid from buffalos, lions, wildebeest, impala, zebra and hyena that were collected in relation to different cross-sectional studies conducted in the Serengeti ecosystem in Tanzania were used. Multiplex polymerase chain reaction AMOS-PCR and quantitative Real-Time PCR (qPCR) targeting the genus specific surface protein bcsp31 gene and the insertion sequence IS*711* element downstream of the *alkB* gene for *B. abortus* and *BMEI*1162 gene for *B. melitensis* were employed on the samples.

**Results:** Results indicated that out of 189 samples examined, 12 (6.4%) and 22 (11.6%) contained *Brucella* DNA as detected by AMOS-PCR and qPCR, respectively. Most of the positive samples were from lions (52.6%) and buffaloes (19.6%). Other animals that were positive included wildebeest, impala, zebra and hyena. Out of 22 positive samples, 16 (66.7%) were identified as *B. abortus* and the rest were *B. melitensis.*

**Conclusion:** Detection of zoonotic *Brucella* species in wildlife suggests that livestock and humans at the interface areas where there is high interaction are at risk of acquiring the infection. Therefore, public education to interrupt risky transmission practices is needed. The findings also shed light on the transmission dynamics around interface areas and the role of wildlife in transmission and maintenance of *Brucella* infection in the region.

## Introduction

Brucellosis is a bacterial zoonosis of public health and economic importance world-wide. It affects a number of livestock and wildlife species including humans [1]. The disease is a challenging public health problem to control in many developing countries including Tanzania, especially in pastoral and agro-pastoral farming systems [2–4]. According to WHO, brucellosis is one of the important re-emerging neglected tropical zoonosis [5] largely due to lack of public awareness.

In wild animals, brucellosis can be a result of spill-over from infected livestock or as a natural sustainable infection within susceptible wild animal populations [6,7]. Wild ungulates could acquire infection by ingesting contaminated pasture [7]. Flesh-eaters such as wolves and foxes are thought to be exposed through the ingestion of infected animals, placentae or aborted foetuses [8]. The disease has been reported in wild animals in some African countries, which include Kenya [9], South Africa [7], Zimbabwe [10] and Tanzania [11–14]. In the later country wild animals, *Brucella* infections have been reported in topi, buffalo, impala, Thompson gazelle and wildebeest [15,16]. However, most of these studies have been serology based without indication of circulating *Brucella* spp. Other studies reported brucellosis in livestock-wildlife interfaces in the Ngorongoro Conservation Area and Mikumi Selous Ecosystem [12–14].

In recent years many African countries have prioritized zoonotic diseases under the global Health Security Agenda. In many countries to date, brucellosis has been ranked among important zoonoses. In Tanzania in particular, it ranks among the top six priority zoonoses that the country will focus control efforts [17]. Since the prioritization of brucellosis in 2017, there have been efforts for development of a control strategy. In implementation of that strategy, critically highlighted areas include the pattern and contribution of different hosts in the transmission and maintenance of brucellosis in the country. Studies have been done on the livestock and shed light on *Brucella* species circulating in different regions of the country [18,19]. However, wildlife transmission dynamics remain a grey area. The aim of this study was to determine circulating *Brucella* species in wildlife in Serengeti ecosystem in Tanzania, using molecular techniques.

## Materials and methods

### Study area

The study used samples of cross-sectional studies conducted so far in Serengeti ecosystem in Tanzania. The Serengeti is the world’s most diverse ecosystem, located in the in north-west of the country between the Ngorongoro highlands and Lake Victoria. This ecosystem comprises of Serengeti National Park, the Ngorongoro Conservation Area, Maswa Game Reserve, Loliondo Game Controlled Area and Kenya’s Masai Mara National Reserve (Fig. 1). The study area was selected because there is notable interaction between wild animals, livestock and humans. This area is mainly inhabited by the Maasai with livestock keeping being their main economic activity [20].

**Figure 1:**
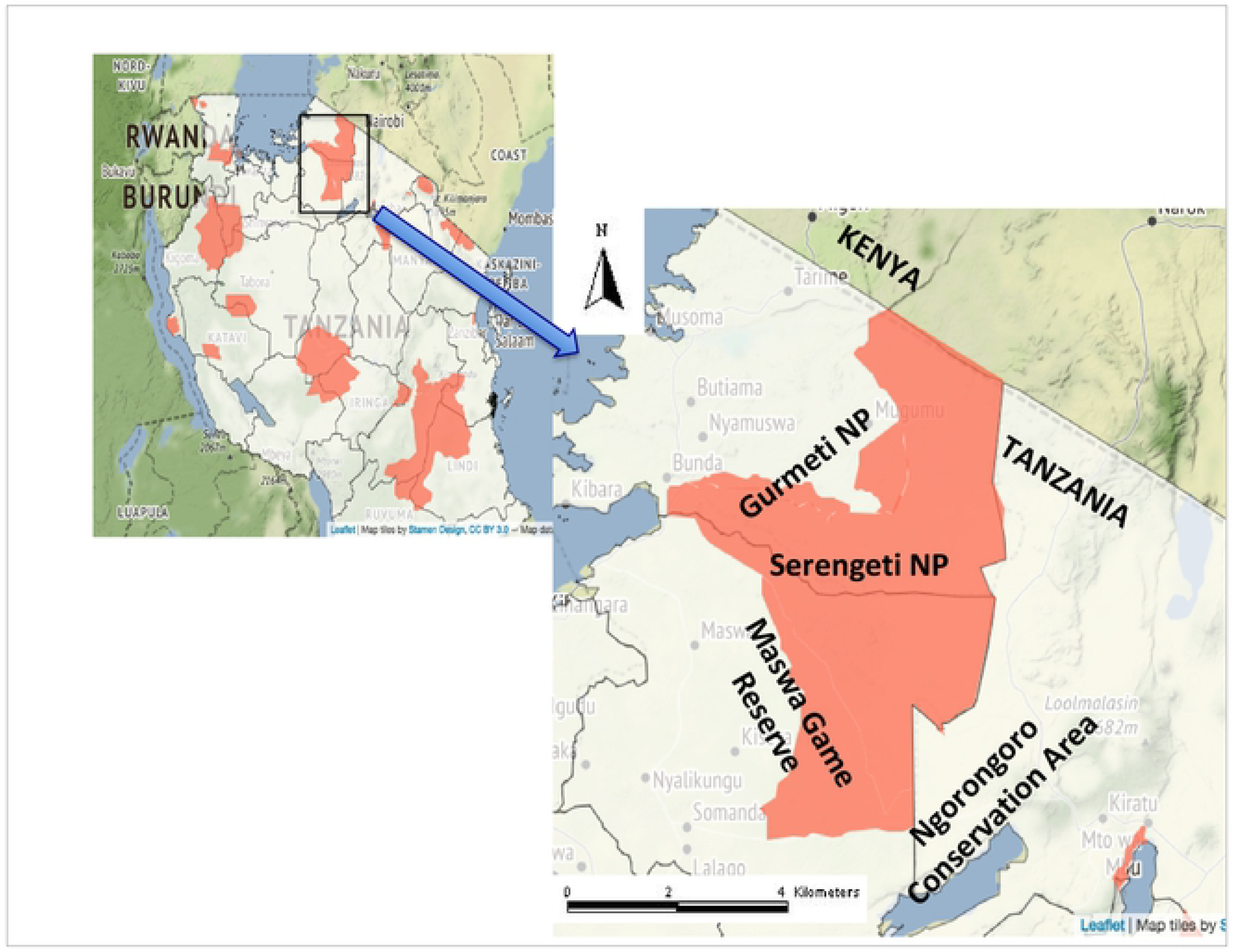
A map of the study area showing Tanzania conservation areas and the Serengeti ecosystem

ecosystem, National Park and the Ngorongoro Conservation Area (expanded insert). Samples used in this study were collected between 2010 and 2017 during routine surveillance and veterinary training programs

### Collection of biological samples

Serum, whole blood and amniotic fluid samples were used in the present study. The samples were collected in connection to other research activities between 2000 and 2017 and archived in the Tanzania Wildlife Research Institute (TAWIRI) biorepository in Arusha, Tanzania and Serengeti laboratory and stored at -20°C. The samples were collected from buffaloes, wildebeest, zebra, lions, baboon, impala and hyena. In total, 189 samples were collected out of which 11 were amniotic fluid, 170 whole blood and 8 serum samples.

### Molecular detection of *Brucella* spp

The study employed AMOS PCR and a quantitative Real-Time PCR (qPCR) in the detection of *Brucella* spp. from the samples. The PCR protocols used were as described elsewhere [21–23]. In detail, at the Microbiology labs, college of veterinary medicine and biosciences in Sokoine University of Agriculture (SUA) Tanzania, samples were subjected to DNA extraction using a commercial DNA extraction kit (Zymo Research, USA Genomic DNA™ Tissue Mini Prep kit) and a method as described by Navarro *et al.* (2002) [24]. Briefly, 40µl of genomic lysis buffer was added to 200µl of the source sample. The mixture was subjected to digestion, deactivation, washing and elution steps as per manufacturer’s instructions. Stock DNA samples were stored at -20 °C until the performance of PCR. Primers used in this analysis were obtained from Bioline Inc (Taunton, MA, USA) and are detailed in Table 1.

**Table 1:**
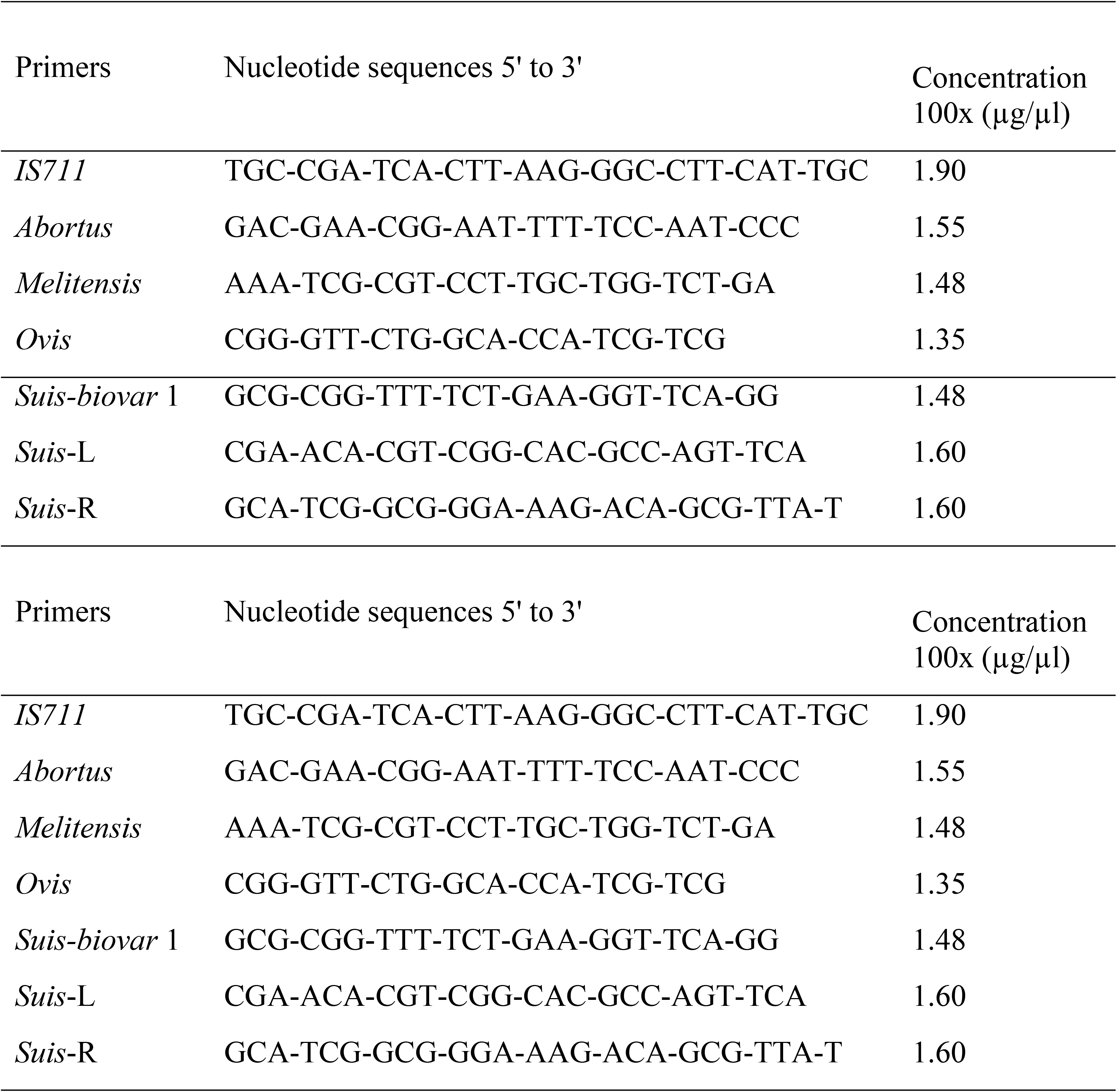
Primers used to amplify the target region of *Brucella* spp. present in the DNA extracts using the AMOS PCR.

The quantitative qPCR for *Brucella* spp. identification targeting the *bcsp31* gene (GenBank accession number M20404) and *IS711* (GenBank accession number HE603358.1) genes were used [21,23]. Samples positive for the *Brucella* genus level target were then subjected to a multiplex assay to distinguish *B. abortus* from *melitensis*. The assay used *B. abortus* and *B. melitensis* primers targeting the specific insertion of an IS*711* element downstream of the *alkB* (GenBank accession number AF148682) and *BMEI1162* (access number NC_003317) genes respectively. Analysis was done according to manufacturer instruction in the *Brucella* genus Genesig® standard kit (Genesig® Camberly, UK). A volume of 10µl DNA was mixed with primers and probes in 1000µl reaction tubes as described elsewhere [23]. Primers and probes used in the qPCR assay for the detection of *Brucella* spp. are shown in Table 2. Amplification and real-time fluorescence detection were performed on the iCycler real-time PCR detection system (Bio-Rad Laboratories, Hercules, Calif.). Positivity criteria of the assay required that a sample amplifies in both targets and below a set amplification cycle time (<38).

**Table 2:**
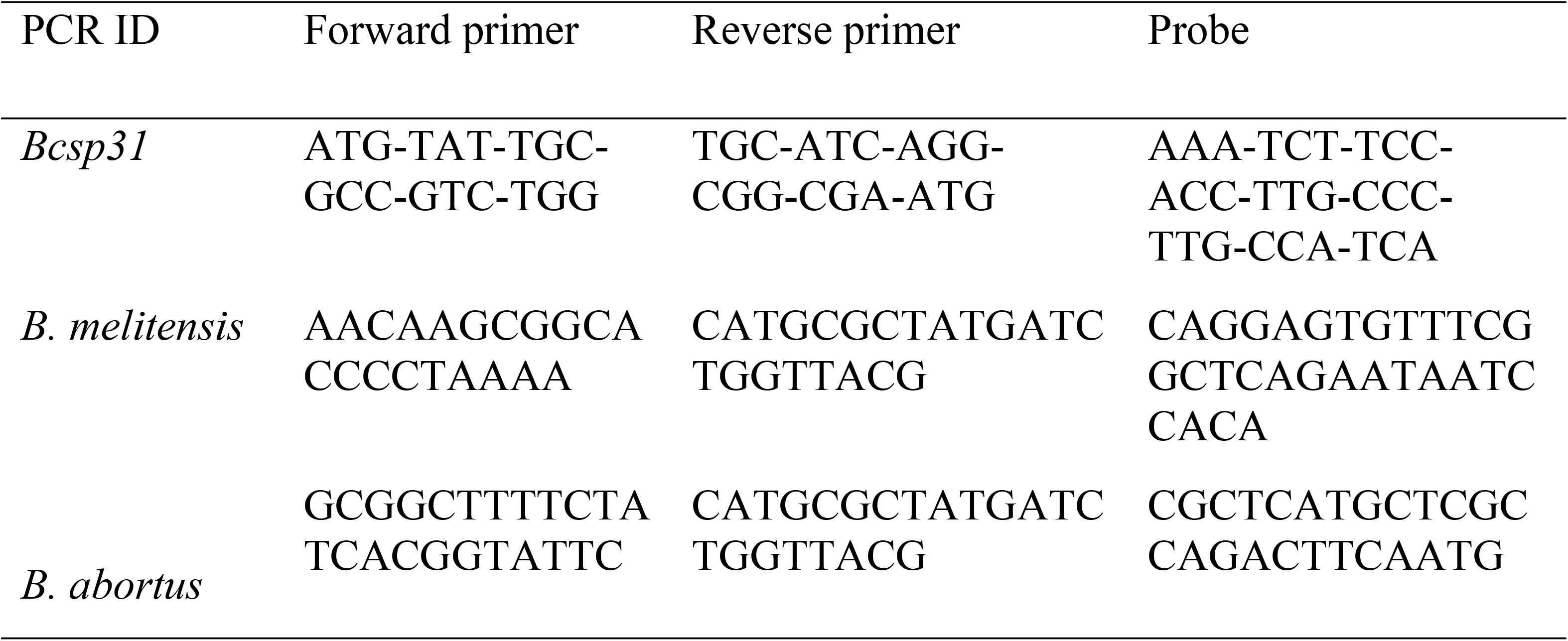
Primers and probes used in the real-time multiplex PCR assay for the detection and speciation of *Brucella* spp.

The results from each of the techniques were then pooled and cleaned in Microsoft Excel® (IBM, USA v. 2010) then descriptive and crude analytical statistics done using R software (25). Chi-square (χ^2^) or Fisher’s exact test were used as appropriate for comparison of; age, sex, location, animal species and *Brucella* PCR positivity were done. Population differences with a p value < 0.05 were considered as significant. Cross tabulation was used to determine the diagnostic sensitivity and specificity of the AMOS and real-time qPCR using the qPCR speciation assay as the reference test.

### Ethical consideration

This study was conducted in conformity with the ethical and animal welfare guidelines stipulated by Sokoine University of Agriculture research ethics. A research permit was provided by the Tanzania Wildlife Research Institute (TAWIRI) for study conduct in wildlife (TWRI/RS/57/VOL IV/85/72).

## Results

A total of 189 samples from seven wild animal species, (buffaloes, wildebeest, zebra, lions, baboons, impala and hyenas) were used in the present study. Out of these, 170 whole blood collected in the EDTA tubes, 8 sera and 11 amniotic fluid samples.

Most of the samples (80; 42.3%) were from wildebeest. Most samples (183; 96.8%) used were obtained from female animals. In terms of specific location, the majority (115; 60.9%) of the samples were from Serengeti National Park. On crude association with study variables, it was found that the age of the wild animal sampled (Adult), location (Serengeti) and the sample type used for DNA extraction (EDTA whole blood) were all significantly associated with the detection of *Brucella* DNA (Table 3).

**Table 3:** Characteristic features of the samples used in the study

| Variable | Categories | Number (%) | Pearson's $\chi^2$ | p-value |
| --- | --- | --- | --- | --- |
| Sex | Female | 183 (96.80) | 1.08 | 0.30<sup>a</sup> |
|  | Male | 6 (3.20) |  |  |
| Age (group) | Adult | 186 (98.41) | 4.36 | **0.04<sup>b</sup>** |
|  | Sub-adult | 3 (1.59) |  |  |
| Location | Serengeti | 115 (60.85) | 0.27 | 0.60<sup>a</sup> |
|  | Ngorongoro | 74 (39.15) |  |  |
| Species | Buffalo | 46 (24.34) | 23.20<sup>b</sup> | **0.001<sup>b</sup>** |
|  | Wildebeest | 80 (42.33) |  |  |
|  | Zebra | 25 (13.23) |  |  |
|  | Lion | 19 (10.05) |  |  |
|  | Baboon | 5 (2.65) |  |  |
|  | Impala | 10 (5.29) |  |  |
|  | Hyena | 4 (2.12) |  |  |
| Sample type | EDTA | 170 (89.95) |  | **0.04<sup>a</sup>** |
|  | Serum | 8 (4.23) | 6.21 |  |
|  | Amniotic fluid | 11(5.82) |  |  |

N= number of individuals in each category. This data stem from the study conducted between 2010 and 2017 in the Serengeti National Park and Ngorongoro Conservation area.
<sup>a</sup> P-value based on $\chi^2$ -test; <sup>b</sup> P-value based on Fisher's exact test; Statistically significant ( $p<0.05$ ) are highlighted in bold

Of the 189 samples screened, *Brucella* DNA were extracted from 12 (6.4%) samples. Based on AMOS PCR, 16 (66.7%) out of the 24 positive samples were identified as *B. abortus*, 2 (8%) were *B. suis* and 2 (8%) *B. melitensis*. The animal species distribution of *Brucella* DNA positive samples based on AMOS PCR are detailed in Table 4. Two samples; from buffalo and impala were found to harbor more than one *Brucella* species each. The species that were found in a single animal included *B. melitensis, B. suis* and *B. abortus* suggesting multiple infections.

**Table 4:** *Brucella* spp. in wild animals in Serengeti ecosystem detected by AMOS-PCR and qPCR (n=189)

| Animal spp. | AMOS PCR | q PCR |
| --- | --- | --- |
| Lion | *Brucella abortus* (n=3) | *B. abortus* (n=6) |
| Buffalo | *Brucella abortus* (n=1) | *B. abortus* (n=7) |
| Wildebeest | *Brucella suis* (n=4) | NA |
| Zebra | *Brucella suis* (n=1) | *B. abortus* (n=1) |
| Impala | *Brucella melitensis* (n=1) | *B. abortus* (n=2) |
| Hyena | *Brucella suis* (n=1) | NA |
| Baboon | NA | NA |

NA= No amplification. This data stem from a study conducted between 2010 and 2017 in the Serengeti National park and The Ngorongoro Conservation area

The qPCR test results indicated that 22 samples (11.6%) were positive for *Brucella* DNA. Overall, 16 samples out of 22 (66.7%) samples were positive for *B. abortus* in qPCR. The same 16 samples were also positive *for B. abortus* in AMOS PCR. Using the real-time speciation assay as the reference test, AMOS PCR had a sensitivity of 16.7% and specificity of 92%, while the qPCR assay had a sensitivity of 72.7% and specificity of 100% (Table 5)

**Table 5:** Cross tabulation of the molecular tests used, with RT-qPCR speciation assay as the reference

|  | **RT-Speciation** |  |  |
| --- | --- | --- | --- |
|  | **Positive** | **Negative** | **Total** |
| **AMOS-PCR** |  |  |  |
| Positive | 2 (0.167) | 10 (0.053) | 12 (0.063) |
| Negative | 14 (0.074) | 163 (0.921) | 173 (0.937) |
| **RT-qPCR** |  |  |  |
| Positive | 16 (0.727) | 6 (0.032) | 22 (0.116) |
| Negative | 0 (0) | 167 (1.000) | 167 (0.884) |
| **Total** | 16 (0.085) | 173 (0.915) | 189 (100) |

## Discussion

*Brucella* spp. were detected in wildlife in the Serengeti ecosystem. More *Brucella* spp. DNA were detected in the screened samples using qPCR than AMOS PCR. Lions and buffaloes had the highest proportions of positivity from the sample pool. The most commonly identified species in the wild animals *was Brucella abortus* although *B. suis, and B. melitensis* were also detected. This is the first reported study to conduct molecular detection of *Brucella* directly from archived samples.

Results obtained from qPCR show that *B. abortus* was dominant in the samples collected suggesting that it is a common *Brucella* species circulating in Serengeti ecosystem. Detection of *Brucella* spp. from the study area is not surprising as previous studies have reported *Brucella* sero-positivity in wild animals in Tanzania including Serengeti with ranges between 10.5% and 17% [12,15,16,18,25]. Therefore, detection of pathogenic DNA in samples collected from wildlife in the area confirm that *Brucella* is circulating in the studied ecosystem.

Most of the *Brucella* positive samples were detected in female animals 30 (16.4%) different from the previous report in Tanzania [18] and elsewhere in Africa (3). The current finding further stresses the role of female animals in the transmission and maintenance of *Brucella* infection. During abortion or normal birth there is massive shedding of *Brucellaceae* in the environment which are likely to be picked by susceptible animals during grazing [20].

It was further observed that *Brucella* DNA were detected more in lions (52.6%) and buffaloes (19.6%) than in other wild animal species. The findings could probably be because lions are indiscriminate carnivores and are likely to prey on *Brucella* infected animals like buffaloes. However, the high detection rates observed in buffaloes may be due to *B. abortus* being the common species in the ecosystem and is known to mostly infect bovine ungulates. Generally, detection of zoonotic *Brucella* in wildlife in this study, point to their possible involvement as the source of sustained *Brucella* transmission in livestock and humans in the interface areas of Serengeti ecosystem. It has been earlier reported that wildlife can act as a source of infection for livestock and humans [25–27].

It was also observed that there was higher *Brucella* infection in Serengeti National Park than in the Ngorongoro Conservation Areas probably because Serengeti is a niche habitat of lions and buffaloes and hence contamination of environment is likely to be high [13]. In addition, in the Ngorongoro conservation area there are close interactions between livestock and wild animals [28]. However, it is unclear whether sustained *Brucella* infection in wildlife is acquired from livestock or vice versa. Nevertheless, vaccination in livestock may minimize the reverse spread of the disease from livestock to wildlife and vice versa [28].

Wildebeest seasonally migrate from Serengeti to Masai Mara for pastures [29], a practice likely to spread *Brucella* in the Serengeti ecosystem. Zebra constantly intermingle with wildebeest during grazing [29]; living together in close association and this behavior could be the basis for the transmission of the pathogens amongst the wild animals. Detection of *B. suis* in hyena could be explained by the scavenging behavior in this species.

In this study some animals were detected to have more than one of *Brucella* spp. *B. abortus* and *B. melitensis*, and *B. abortus* and *B. suis* were detected in one individual indicating occurrence of multiple infection as it has been reported before [30]. This study also reported the occurrence of *B. melitensis* and *B. abortus* in Impala. Studies have associated high sero-prevalences in antelope in Kafue flat area in Zambia because cattle were sharing source of water with wild animals during dry season [2,27]. Infection of *B. melitensis* is reported to be less common in sub-Saharan African countries [31,33]. *Brucella melitensis* preferably infect sheep and goat which are related with impala [34].

In this study, qPCR was observed to have a higher detection rate of *Brucella* spp. than AMOS PCR. This finding is supported by reports from other studies which reported qPCR as superior tool [21,23,35,36]. Previous studies that have compared the two platforms and reported similar performance in the detection of *Brucella* DNA [21,36]. This could probably be because AMOS PCR is limited in the detection of all *Brucella* spp. biotypes. Depending on which biotypes are predominant in the region, example *B. abortus* biovar 3 which has previously been detected in Tanzania [19] cannot be detected in AMOS PCR. Although the assay sensitively detected Brucella DNA in these archived samples, we did not have sufficient quantities and quality to confirm genomic material harvested from such archived samples.

## Conclusion

Findings from this study show that *Brucella* spp. are circulating in different wildlife spp. in Serengeti ecosystem. The molecular assays run used DNA extracted from archived samples, indicating their potential for routine surveillance of brucellosis on clinical samples. Most of *Brucella* spp. detected have zoonotic potential. Detection of zoonotic *Brucella* species in wildlife suggests that livestock and humans at the interface areas are at risk of acquiring the infection. The findings also shed lights on the transmission dynamics around interface areas and the role of wildlife in maintenance and transmission of *Brucella* infection in the region. The findings from this study, although contextual to the Serengeti ecosystem, provide valuable insights into *Brucella* infection and host associations in wildlife applicable to much of sub-Saharan Africa

## Conflict of interest

The author declares no conflict of interest. The findings and conclusions in this paper are those of the authors and do not necessarily represent the official position of the participating institutions or the funding organization.

### Acknowledgements

SRM, CM, AL and RBY acknowledge support from the DELTAS Africa Initiative [Afrique One-ASPIRE /DEL-15-008]. Afrique One-ASPIRE is funded by a consortium of donor including the African Academy of Sciences (AAS) Alliance for Accelerating Excellence in Science in Africa (AESA), the New Partnership for Africa’s Development Planning and Coordinating (NEPAD) Agency, the Wellcome Trust [107753/A/15/Z] and the UK government.

Laboratory technicians at Sokoine University of Agriculture, and College of Veterinary of Medicine and Biomedical Sciences and Kilimanjaro Clinical Research Institute-Biotechnology Laboratories for technical assistance, Tanzania Wildlife Research Institute (TAWIRI) for their willingness in sharing the samples.

## Author contributions

**Concept development;** Bonfoh B., Kazwala R. R, Fokou G.

**Funding acquisition;** Bonfoh B., Kazwala R. R, Fokou G.

**Investigation and Formal analysis:** Sambu, R. M, Nonga, H.E., Mathew C., Lukambagire A. S.,

**Methodology;** Sambu, R. M, Mathew, C., Lukambagire A. S., Richard B. Yapi, Kazwala R. R

**Supervision;** Nonga, H.E., Mathew C, Keyyu J., Kazwala R. R:

**Writing – original draft;** Sambu, R. M., Mathew C.

**Writing – review & editing**; All Authors

